# DNA-Demethylating Agents enhance cytolytic activity of CD8+ T Cells and anti-tumor immunity

**DOI:** 10.1101/197236

**Authors:** Helen Loo Yau, Ankur Chakravarthy, Felipe Campos de Almeida, David Allard, Rajat Singhania, Ilias Ettayebi, Shu Yi Shen, Tiago Medina, Parinaz Mehdipour, Beatriz Morancho, Sandra Pommey, Christian Klein, Gustavo Amarante-Mendes, David Roulois, Marcus Butler, Joaquín Arribas, John Stagg, Daniel D. De Carvalho

## Abstract

Recent studies have shown that DNA methyltransferase inhibitors (DNMTi) can induce IRF7 activation and Type I/III interferon signaling through dsRNA-mediated viral mimicry in cancer cells. By performing a large pan-cancer analysis using TCGA data, we determined that IRF7 activation is associated with higher CD8+ T cell tumor infiltration and higher cytolytic activity across multiple cancer types. Accordingly, we demonstrate that DNMTi treatment results in increased CD8+ T cell tumor infiltration, enhanced cytolytic activity and CD8+ T cell dependent tumor growth inhibition. Finally, we show that DNMTi triggers a process marked by the induction of viral mimicry directly on CD8+ T cells, leading to activation of dsRNA sensing pathway, and up-regulation of T cell activation markers, effector cytokines, and Granzyme B. Taken together, our findings suggest that dsRNA sensing pathway activation in the immune compartment, through pharmacological DNA demethylation, is a viable strategy for boosting anti-tumor immune response.

## Introduction

The use of DNA methyltransferase inhibitors (DNMTi) such as 5-Azacytidine and Decitabine (5-AZA-CdR) has recently emerged as attractive therapeutic alternatives for solid cancers. Most recently, these DNA demethylating agents were shown to have long-term effects, including the depletion of the stem cell compartment of colorectal cancers^1^. This effect was shown to be mediated through the induction of a state of viral mimicry, induced by global DNA demethylation leading to the transcriptional upregulation of human endogenous retroviruses (HERVs), dsRNA formation, activation of the dsRNA sensing pathway MDA5/MAVS/IRF7, and ultimately, the production of Type III Interferon^1^. Moreover, a signature of Interferon Response Genes attributed to viral mimicry, was linked to differences in responses to immune checkpoint inhibition in patient samples from clinical trials, as well as in pre-clinical murine models of melanoma^2^. These discoveries have fueled interest in the combination of DNMTi, to induce viral mimicry and increase anti-tumor visibility by the immune system, with immune checkpoint blockade^3-5^. However, little is known on the effects of DNMTi on the immune compartment, especially at clinically relevant doses.

Since DNMTi can modulate Type I and III Interferon^1,2^ signaling, we sought to contextualize the role of Interferon Response Signaling in tumors with respect to the composition of the immune infiltrate and immune activity within tumors. We also evaluated the effects of the DNMTi-mediated Interferon Response Signaling within the immune compartment. We employed bioinformatics tools that integrated DNA methylation-based estimates of immune cell populations with transcriptomics-based measurements of cytolytic activity and neoantigen burden. From these estimates, we demonstrate that an IRF7 activation signal is independently associated with higher CD8+ T cell infiltration and higher cytolytic activity in multiple tumor types. Remarkably, this higher IRF7 signal seems to originate mainly from the immune infiltrate of these tumors. Indeed, *in vivo* and *ex vivo* experiments of syngeneic tumor models and human CD8+ T cells treated with clinically relevant doses of DNMTi highlights the ability of DNA demethylating drugs to activate the same IRF7-mediated transcriptional patterns associated with an active immune infiltrate phenotype. DNMTi treatment leads to enhanced CD8+ T cell infiltration and enhanced CD8+ T cell effector function associated with enhanced cytolytic activity. Moreover, depletion of CD8+ T cells is sufficient to abolish the anti-tumor effect of DNMTi treatment *in vivo*. Finally, we show that clinically relevant doses of DNMTi can activate a viral mimicry phenotype characterized by increased expression of HERVs, activation of the dsRNA sensing pathway and IRF7 activation directly on human CD8+ T cells. This IRF7 activation is directly associated with enhanced expression of the effector molecules interferon gamma (IFN-ɣ) and Granzyme B by the CD8+ T cells. Altogether, our results highlight novel immune-specific immunomodulatory properties of DNMTis.

## Results

### Activation of interferon signalling is associated with distinct patterns of immune infiltration

We previously showed that the viral mimicry state, induced by treatment with DNMTi, involved the downstream activation of IRF7^1^. This IRF7-dependent viral mimicry signature is associated with better response to immunotherapy^2^. Therefore, we asked if we could observe distinct immune cell infiltration patterns between tumors stratified by IRF7 signalling. To achieve this, we generated immune cell estimates using MethylCIBERSORT^6^ for more than 7,000 tumors across 19 distinct tumor types with data obtained from The Cancer Genome Atlas (TCGA). Following quality control and integration with matched expression data, we derived a set of 5,996 tumors. To partition samples based on IRF7 activation, we used PAM clustering to search for the optimal number of clusters and identified two broad clusters: IRF7-high (C1) and IRF7-low (C2) (Figure 1A). Integrating this with cellular estimates, we found the vast majority of infiltrating cell types to be differentially abundant between the clusters (Figure 1B). Remarkably, there was a large increase in CD8+ T cell infiltration in tumors from the IRF7-high cluster, suggesting a potentially greater immune activity. This finding is relevant as CD8+ T cell tumor infiltration is a known predictor of response to anti-PD1 checkpoint therapy^7^, as well as prognosis in multiple cancers^8-10^. We also observed some increase in Tregs in tumors from the IRF7-high cluster (Figure 1B), probably an indication of selective pressures to suppress cytolysis by CD8+ effectors^11^

**Figure 1.**
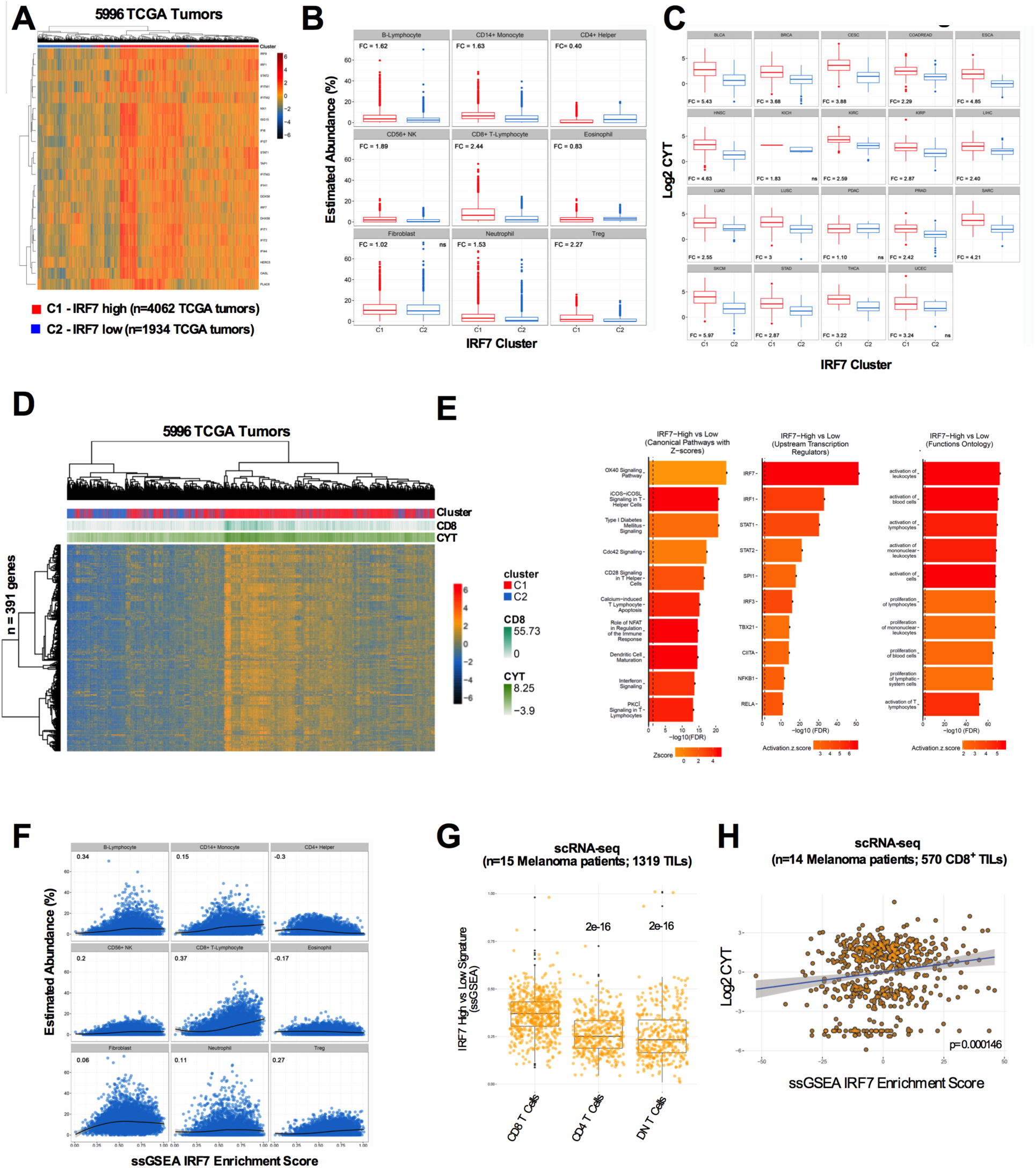
IRF7-high tumors have increased immune-infiltration associated with high CD8+ T cell activation and cytolytic activity. **A)** Heatmap showing the expression profile of IRF7 “viral mimicry” signature genes in 5,996 TCGA RNA-seq data from multiple cancer types. **B)** MethylCIBERSORT estimates show that IRF7-High tumors (C1) are associated with distinct patterns of cellular infiltration. X-axis = C1, IRF7-high cluster; C2, IRF7-low cluster. Y-axis = estimated abundance of cell type. FC = mean fold change, Plots with ‘ns’ are not significant, all the other plots are significant at FDR < 0.01. FDR from Wilcoxon’s Rank Sum Tests. **C)** Boxplots of cytolytic activity by IRF7-cluster. Cytolytic activity was quantified as in Rooney et al^11^. X-axis = IRF7 cluster, Y-axis = log2(Cytolytic Activity). Plots with ‘ns’ are not significant; all the other plots are significant at FDR < 0.01. FDR from Wilcoxon’s Rank Sum Tests. **D)** Heatmap of genes differentially expressed between the two IRF7-Signature clusters. **E)** Results of IPA analyses of canonical pathways, upstream regulators and functional ontology (Top 10 pathways with available activity patterns in each case) performed on genes differentially expressed between IRF7-high and IRF7-low tumors. Intensities represent activation Z-scores, while X axes represent -log10(FDR). **F)** Correlation scatterplots of IRF7-ssGSEA (X-axis) scores versus estimated abundances of each infiltrating cell type (Y-axis); values represent Spearman’s Rho. **G)** Boxplots of the extended 391 gene IRF7 signature ssGSEA scores (Y-axis) in single cell RNA-seq (scRNA-seq) data from CD8+ T cells, CD4 T cells and double negative T cells from 1319 TILs obtained from 15 Melanoma patients. scRNA-seq data was obtained from Tirosh et al^15^. FDR from Wilcoxon’s Rank Sum Tests versus CD8+ T Cells. **H)** Added-variable scatterplot scatterplots of patient-corrected IRF7-ssGSEA (X-axis) scores versus cytolytic activity (Y-axis) on 570 CD8+ TILs. scRNA-seq data was obtained from Tirosh et al^15^.

Across the vast majority of tumor types, we confirmed that IRF7-high tumors demonstrate significantly elevated cytolytic activity (CYT) (Figure 1C), a surrogate for immune activity with predictive ability in determining immunotherapy response^11^.

Subsequently, regressing log2(CYT) versus infiltrating cell estimates and IRF7-cluster as predictors demonstrated that IRF7-cluster was an independent predictor of cytolytic activity (IRF7-low estimate -0.86, p-value < 2.2e-16). This suggests that IRF7 activation is representative of elevated cytolytic activity even after controlling for the different abundances of CD8+ TILs by cluster and across tumors, implicating that CD8+ T cells with higher IRF7 activation may have greater cytolytic activity and killing potential.

### IRF7-high tumors display a distinct transcriptional profile consistent with greater CD8+T cell infiltration and cytolytic activity

To further probe the transcriptional correlates of IRF7-associated clusters, we used *limma-trend* analysis and discovered large-scale transcriptional differences between the two IRF7-clusters after controlling for cancer type, identifying 391 Differentially Expressed Genes (DEGs) (Figure 1D) (2FC, FDR < 0.001, Supplementary Table S1). Ingenuity Pathway Analysis (IPA, www.ingenuity.com) correctly inferred the activation of IRF7 and identified a large set of lymphocyte maturation and activity related genes up-regulated in the IRF7-high cluster (Figure 1E and Supplementary Tables S2, S3, and S4). Taken together, the expression data corroborates the DNA methylation data, suggesting IRF7-high tumors have an increased and more activated immune infiltrate, suggesting a potentially greater immune activity.

Given that neoantigen burden has been postulated to be a predictor of infiltration and activity of T cells^11,12^, we tested for differences between the clusters in the burden of neoantigens per tumors for 3999 tumors from 17 cancer types. Mainly, we tested if IRF7-clusters were marked by elevated neoantigen burdens. When comparing different tumor types, we found no significant differences between IRF7-high and low tumors in any individual tumor types at FDR < 0.05 (Wilcoxon’s Rank Sum Test), suggesting that IRF7 activation is not simply a feature of neoantigen-dense tumors, consistent with published data showing decoupling of a T cell inflamed transcriptional signature from neoantigen density in melanomas^13^

### Statistical inference suggests the immune compartment is the origin of the IRF7 activation signal in tumors

In order to infer the likely source of the IRF7 activation signal, we initially characterised the type of interferon response associated with IRF7 activation using linear regression comparing ssGSEA enrichment scores^14^ for Type I, II and III interferons against an ssGSEA score for IRF7 activation, and accordingly found that all three classes of interferon were significant predictors of the IRF7 enrichment score (Figure S1A), with Type III and Type I interferons demonstrating the strongest association per unit increase even if Type II interferons, hinting at a Tumor Infiltrating lymphocyte (TIL) based origin, were the most differentially expressed (2FC, FDR < 0.01).

To further resolve the source of the IRF7 signal, we compared tumor purity and IRF7 ssGSEA scores and found strong inverse correlations across the majority of tumor types (Figure S1B). Subsequently, we also found significant positive correlations between estimated abundances of several immune cell subsets and IRF7-activation ssGSEA scores, with CD8+ T cells showing the strongest correlation (Figure 1F). Then, we used multivariate linear modeling and found multiple immune cell types, including CD8+ T cells, to be a significant predictor of the IRF7 ssGSEA score in the joint analysis, further implicating the immune infiltrate as the source of the IRF7 signal (Table S5).

Then, we tested the hypothesis that CD8+ TILs were a key source of the IRF7 signal in single cell RNA-seq data from dissociated TILs in melanoma patients^15^. We found markedly greater enrichment of the 391-DEG IRF7-associated signature in CD8+ cells (Figure 1G) relative to CD4+ and double-negative TILs. We also found that IRF7 ssGSEA scores were a significant predictor of CYT as predicted from bulk data (Figure 1H) by linear modelling after adjusting for patient (coefficient = 0.024, p = 0.000146) and in univariate analyses based on quartiles of IRF7 ssGSEA scores (Upper vs lower quartile p value = 0.00021, Wilcoxon’s rank sum test, Figure S1C-E).

Finally, to confirm that IRF7-target genes were indeed likely to be transcribed preferentially in the immune compartment of tumors, we obtained publicly available gene expression datasets of colorectal cancers that were sorted into distinct compartments^16^ (GEO accession: GSE39397). Validating our inference from the deconvolution approach, we observed increased IRF7 signature expression within the leukocyte fraction as compared to the tumor epithelial compartment and significantly higher ssGSEA scores in the leukocyte compartments relative to matched epithelial tumor cells (Figure S1F-H).

Taken together, these findings suggest that a higher IRF7 activation in the immune compartment, which is especially enriched for CD8+ T cells, is associated with a greater anti-tumor immune activity, and potentially increased sensitivity to immune checkpoint inhibitors^2^.Consequently, we hypothesized that pharmacologically modulating IRF7 signalling with DNMTi, which we recently showed to be an inducer of IRF7 activation in cancer cells^1^, would produce higher CD8+ infiltration into tumors with corresponding anti-tumor activity and enhance the activity of T-cells by modulating IRF7-associated transcriptional programs through viral mimicry.

### Anti-tumor effect of DNMTi treatment is dependent of CD8+ T cells *in vivo*

In order to experimentally test the hypothesis that DNMTi treatment will modulate CD8+ T cell-dependent anti-tumor immune response, we used the CT26 syngeneic colorectal tumor model^17^. We injected 5x10^5^ CT26 cells in three groups of mice (n=10 mice per group). In the first group, we mock treated. In the second group, we treated with 0.5mg/kg of Decitabine from Day 5 post-tumor inoculation for five consecutive days. In the third group we performed the same Decitabine treatment plus CD8+ T cell-depletion with an anti-CD8+ neutralizing antibody at Days 4, 5 and 10 post-tumor inoculation (Figure 2A). DNMTi treatment significantly reduced tumor growth *in vivo* in CD8+ competent mice, but this anti-tumor effect was abolished in the absence of CD8+ T cells (Figure 2B and Figure S2A).

**Figure 2.**
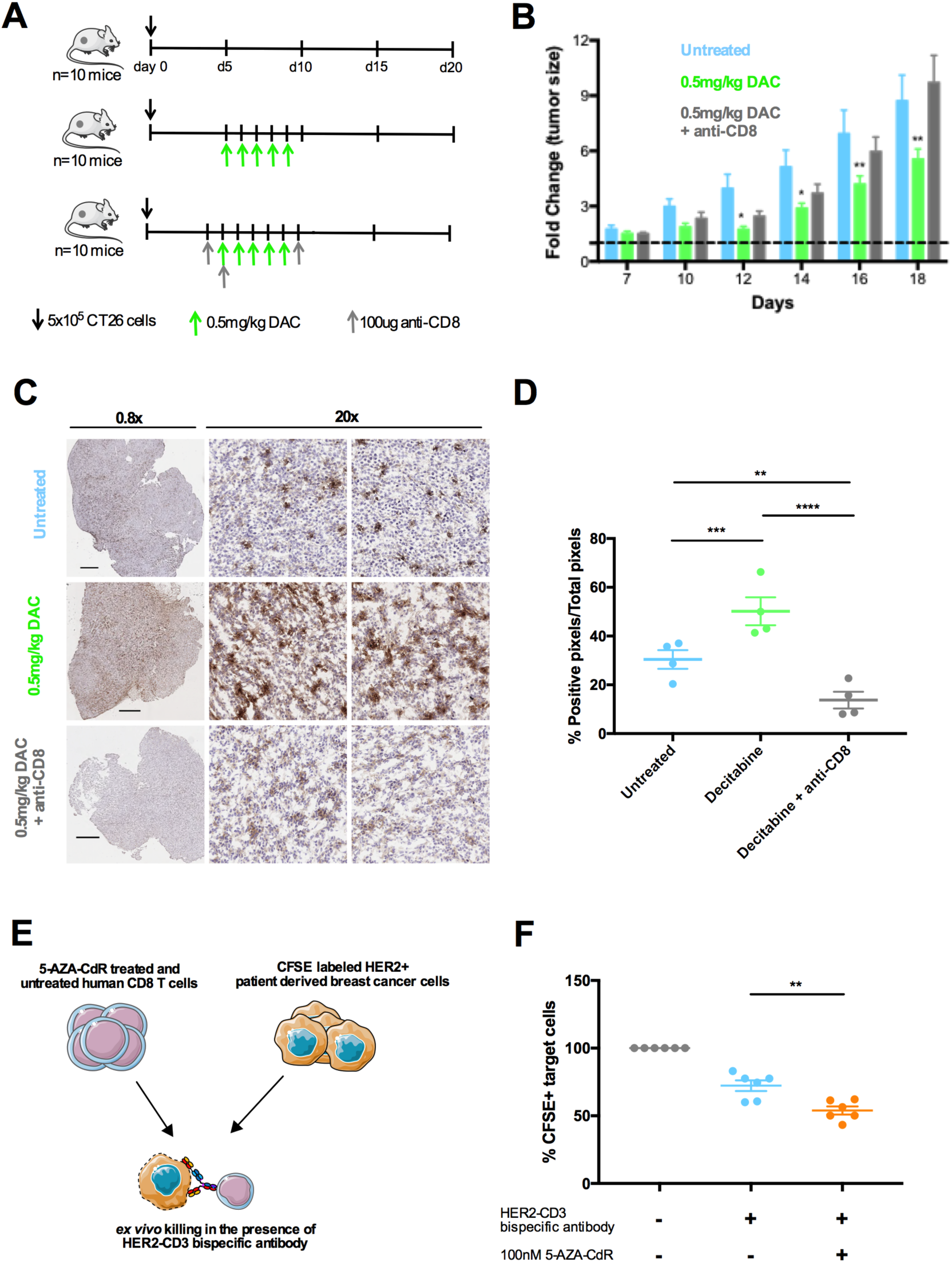
– DNMTi treatment increases CD8+ T cell tumor infiltration and increases CD8+ T cell cytolytic activity. **A)** Schematic representation of the DNMTi treatment. 5×10^5^ cells of the CT26 syngeneic murine cell line were injected subcutaneously into WT BALBc/J mice (n=10 per group) at Day 0. Group one was left untreated. Group 2 was treated with 0.5mg/kg of 5-AZA-CdR at Days 5 to 9. Group 3 was treated with the same dose and schedule of 5-AZA-CdR plus 100ug of neutralizing anti-CD8+ antibody at Days 4, 5, and 10. **B)** Fold change in tumor size measured at Days 7, 10, 12, 14, 16, and 18 post-tumor injection. * p<0.05, ** p<0.01 versus the untreated group (2-way ANOVA, mean ± SEM). **C)** Anti-CD8+ Immunohistochemistry (IHC) staining of a representative tumor for each group. **D)** Anti-CD8+ IHC quantification of four tumors from each group (paired t-test, two sided, mean ± SEM). **E)** Schematic representation of the *ex vivo* DNMTi treatment and killing assay. Human CD8+ T cells were isolated from healthy PBMC and expanded *ex vivo* with 100nM 5-AZA-CdR for 5 days. 5-AZA-CdR treated or untreated CD8+ T cells were then challenged with CFSE-pulsed HER2 positive breast cancer cells at a 1:5 ratio of target to T cell and treated with anti-HER2-CD3 bispecific antibody to induce target cell killing. **F)** Quantification of surviving target cells after the killing assay. ^**^p<0.01 versus the untreated group (paired t-test, two sided, mean ± SEM).

### DNMTi treatment increases CD8+ T cell tumor infiltration and increases CD8+ T cell cytolytic activity

We then stained for CD8+ tumor infiltration for the three treatment groups (Figure 2A) by immunohistochemistry (IHC). As predicted by our pan-cancer TCGA analysis, DNMTi treatment indeed leads to a profound infiltration of CD8+ T cells into the tumor microenvironment (Figure 2C-D).

Our pan-cancer TCGA analysis also predicted that DNMTi treatment could increase CD8+ T cell cytolytic activity. In order to test this hypothesis using primary human T cells without the issue of TCR specificity, we used a HER2-CD3 bi-specific antibody from Roche. This antibody can bind to HER2 in the target cancer cell and to CD3 on the CD8+ T cells, leading to T cell-mediated killing of the target cancer cell. We activated primary human CD8+ T cells *ex vivo* with DNMTi, and then performed the killing assay against HER2 positive breast cancer samples (Figure 2E). We observed an increased expression of the CD25 activation marker on the CD8+ T cells treated with DNMTi (Figure S2B) and an increased killing ability compared to CD8+ T cells activated without DNMTi (Figure 2F). Moreover, we also repeated this experiment against normal breast cells MCF10A where we artificially overexpressed HER2 (Figure S2C). Again, we observed an increased killing ability of DNMTi treated CD8+ T cells compared to untreated CD8+ T cells (Figure S2D).

Finally, in order to test the ability of DNMTi treatment to increase CD8+ T cell killing in a more physiological experimental condition that depends on the TCR-MHC interaction, we used the OT-I TCR transgenic mice model^18^. In this model, all TCR are specific for OVA. We treated the murine CD8+ T cells *ex vivo* with DNMTi and performed an *ex vivo* killing experiment against OVA-pulsed target cells. Again, we observed an increased killing ability of DNMTi treated T cells (Figure S2E).

Altogether, our results suggests that DNMTi could specifically modulate effector activity in CD8+ cells themselves, transcending the inherent limitation of the *in vivo* experiments wherein the effects of DNMTi on anti-tumor immunity may be mediated through effects on cancer/ cells/ or other immune cells, rather than the CD8+ T cells, confirming our second hypothesis. Then, to test if the aforementioned phenotypes were indeed driven through viral mimicry and IRF7 signaling, we performed integrative analyses using global transcriptome, methylome and targeted assays in human CD8+ T-cells treated *ex vivo* with DNMTi

### DNMTi treatment of CD8+ T cells modulates genes associated with IRF7-transcriptional clusters and recapitulates a viral mimicry phenotype

In order to study the influence of DNMTi on CD8+ T cells on a global level, we performed RNA-sequencing and Reduced Representation Bisulfite Sequencing (RRBS) on human CD8+ T cells from peripheral blood of three independent healthy donors. We performed the experiments on unstimulated or resting CD8+ T cells (Day 0), CD8+ T cells stimulated with anti-CD3 and anti-CD28 *ex vivo* in the presence or absence of 300nM of 5-AZA-CdR (Day 5).

Linear modeling on RNA-seq abundance estimates generated using *Kallisto*^19^ identified 1043 DEGs between DNMTi-treated and untreated stimulated CD8+ T cells (2FC, FDR < 0.05, Supplementary Table S6). Interestingly, this included 7 of 22 IRF7 signature genes initially used to identify the IRF7-high and IRF7-low clusters (Figure S3A) and 59 of 391 genes differentially expressed between the IRF7-high versus low clusters (Figure 3A), representing a significant enrichment (Overlap odds-ratio vs median of null distribution = 2.68, p-value < 9.9e-5, permutation overlap test).

**Figure 3.**
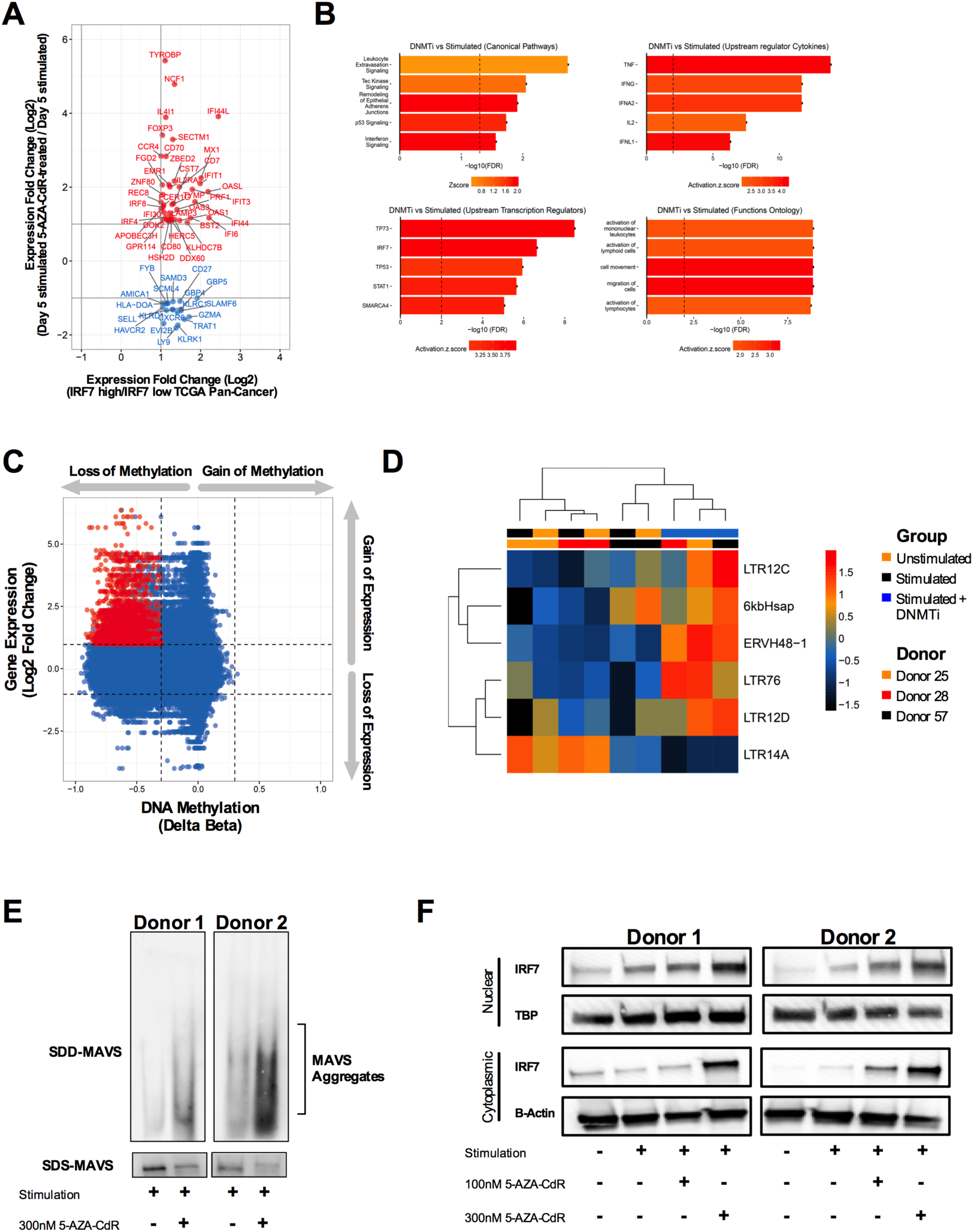
DNMTi treatment leads to viral mimicry, activation of the dsRNA sensing pathway, and IRF7 activation in human CD8+ T cells. **A)** Comparative analysis of fold changes of genes differentially expressed between IRF7-High vs IRF7-Low (2FC, FDR < 0.001) pan-cancer analysis that overlap with genes differentially expressed between human CD8+ T cells activated *ex vivo* with DNMTi treatment vs CD8+ T cells activated without DNMTi treatment (2FC, FDR < 0.05) at Day 5. **B)** Pathway analyses of canonical pathways, upstream regulators and functional ontology (Top 5 pathways with available activity patterns in each case) performed on genes differentially expressed between stimulated CD8+ T cells in the presence or absence of DNMTi. Intensities represent activation Z-scores, while X-axis represents significance (-log10(FDR)). **C)** Quadrant plot plotting magnitude of DNA methylation change (delta-Beta, X-axis) versus magnitude of gene expression (Log2 Fold Change (Y-axis)) of promoter/enhancer/intergenic repeat-associated CpGs. Points coloured red represent MVPs (delta-Beta > 0.3, FDR < 0.01) that are associated with DEGs (logFC > 1, FDR < 0.05) as canonically expected for DNA methylation. **D)** Heatmap showing the ERV-derived transcripts that are differentially expressed (FC > 2, FDR < 0.05) in activated CD8+ T cells in the presence of 300nM 5-AZA-CdR versus activated CD8+ T cells without DNMTi treatment. Rows represent expression Z-scores, columns represent samples. **E)** MAVS aggregation assay. Mitochondrial extracts were isolated from activated (Day 5) human CD8+ T cells from two healthy donors *ex vivo* treated or not with 5-AZA-CdR. The mitochondrial extracts were analyzed by SDD-AGE (top – MAVS aggregation assay) and SDS-AGE (bottom – total MAVS). **F)** Nuclear and cytoplasmic western blots for IRF7 from activated (Day 5) human CD8+ T cells from two healthy donors *ex vivo* treated or not with 5-AZA-CdR.

Pathway analysis on the RNA-seq of DNMTi treated and untreated CD8+ T cells identified two major trends. The first being the upregulation of T cell activation pathways, and the second being the activation of P53 and a DNA damage response. The top canonical pathways included several pathways implicated as critical in lymphocyte-mediated immune responses, including interferon signaling and Tec kinase signaling^20^; the top upstream regulators included multiple cytokines (*TNF, IFNA, IFNB, IFNG, IFNL*), and IRF7 and TP53 amongst upstream transcriptional regulators (Figure 3B, Supplementary Tables S7 and S8). A function ontology analysis identified, amongst others, multiple gene sets associated with lymphocyte activation as being significantly activated (Figure 3B, Supplementary Table S9). This indication of increased activation beyond that seen in stimulated CD8+ T cells was also reflected in the overexpression of multiple key cytokines and effector molecules (*PRF1, GZMB, CCR4, IL15, OX40,* and *4-1BB),* and the down-regulation of the immunosuppressive genes *DGKA* and *HAVCR2 (TIM3).* In mice, *DGKA* has been shown to be required for the induction of anergy^21^. Additionally, TIM3 blockade potentiates immune-mediated tumor clearance by itself, and has the potential for combinatorial use with other immune checkpoint blockade strategies^22^. The down-regulation of these suppressive molecules by DNMTi also constitutes a mechanism for potential synergy with immunotherapy.

In order to understand the role of DNMTi-induced demethylation in the modulation of the IRF7-associated gene set, we carried out RRBS on the samples that were RNA-sequenced, producing Beta-value estimates for 1.7 million CpG sites (> 10X coverage in all samples). Overall, amongst 1,296,010 CpG sites that mapped to regions upstream of genes, FANTOM5 enhancers, or intergenic repeat regions, we identified 330,471 MVPs (methylated variable positions) at absolute Delta-Beta > 0.3, FDR < 0.01, with all MVPs being hypomethylated, confirming that DNMTi induced global hypomethylation (Figure S3B).

Upon integration with 5-AZA-CdR-associated expression profiles, we found 10,450 MVPs associated with 390 DEGs (Figure 3C, Supplementary Table S10), with 2,922 expression-associated MVPs mapping to gene bodies, 222 to FANTOM5 enhancers and 2,106 to promoters. Importantly, there were also 5,200 repeat-associated MVPs in intergenic regions associated with repeat-derived DEGs, with copies of some repeats demethylated on multiple chromosomes (Figure S3C). The most abundantly demethylated (Figure S3C) and re-peats (Figure 3D) repeats were LTR12C and LTR12D, that belong to the Human Endogeous Retrovirus 9 (HERV-9) group. LTR12 elements from HERV-9 are known to be reactivated by DNMTi^2^,23,24 and to have sense and antisense transcription^25^ allowing formation of dsRNA^23^. Moreover, we found that genes with expression-associated MVPs compared to those without were significantly depleted relative to the background distribution of all differentially expressed genes versus non-differentially expressed genes (p-value = 2e-16, proportions test with Yates correction, OR = 0.45), suggesting that direct DNA demethylation plays a relatively small role in the activation of these IRF7 and T cell activation pathways observed in DNMTi treated CD8+ T cells (Figure 3B). The caveat is that demethylation of a small number of candidate regulators may exert a disproportionately large influence on transcriptional programs, even if we did not find evidence for DNA methylation changes at differentially expressed upstream regulators predicted using IPA.

Given that the ability of DNMTi to induce IRF7 activation in cancer cells is dependent on transcriptional activation of endogenous retrovirus^1^,2, we also examined the expression levels of these elements on the CD8+ T cells upon DNMTi treatment. We observed increased expression of several of these sequences (Figure 3D), suggesting that DNMTi could also induce a viral mimicry response in the CD8+ T cells.

To test this hypothesis, we performed a MAVS aggregation assay to monitor activation of the dsRNA sensing pathway^1,26,27^. Indeed, stimulated CD8+ T cells treated with DNMTi show strong MAVS aggregation that is consistent with activation of dsRNA sensing pathway (Figure 3E). Activation of the dsRNA-sensing pathway leads to MAVS aggregation and downstream activation of IRF7 transcription factor. Again, consistent with the activation of dsRNA sensing pathway on CD8+ T cells treated with DNMTi, we observed increase in IRF7 protein levels and increase in IRF7 nuclear translocation in treated cells (Figure 3F).

Altogether, our results suggest that DNMTi-mediated global DNA demethylation in CD8+ T cells leads to DNA demethylation and reactivation of repetitive elements, notably the LTR12 family, activation of the dsRNA sensing pathway, leading to IRF7 activation and up-regulation of IRF7 transcriptional networks (Figure 3). This will ultimately lead to increased T cell killing capacity and anti-tumor immune response (Figure 2). In order to better understand how DNMTi treatment and the downstream IRF7 activation leads to increased T cell killing capacity, we then performed an extensive CD8+ T cell phenotypic characterization upon DNMTi treatment.

### DNMTi treatment during *ex vivo* CD8+ T cell activation leads to decreased expansion and increased cell death

In order to validate the *in silico* predictions made from expression data, we performed a range of quantitative assays *ex vivo*. Polyclonal activation of human CD8+ T cells with anti-CD3 and anti-CD28 induced a massive T cell expansion at Day 5 (Figure 4A-B). Treatment with 5-AZA-CdR led to a decrease in T cell expansion in a dose-dependent manner (Figure 4A-B, p-value < 0.001, paired t-test, two-sided). Moreover, DNMTi treatment induced a small increase in cell death (Figure 4C, p-value < 0.05, paired t-test, two-sided; Figure S4A).

**Figure 4.**
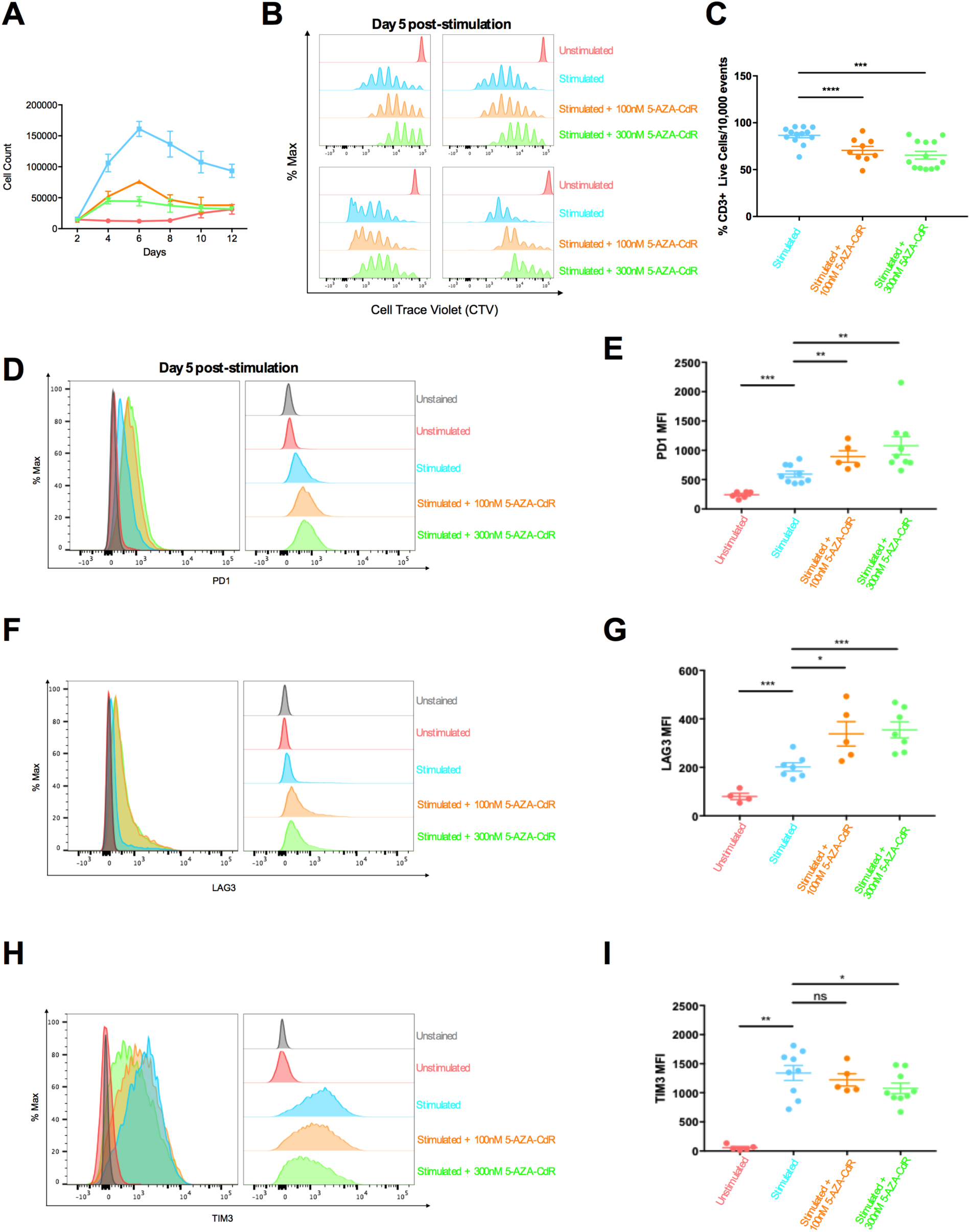
DNMTi treatment delays cell cycle progression, increases cell death and modulates immune checkpoints expression in activated human CD8+ T cells. **A)** Cell count of CD8+ T cells after stimulation with anti-CD3/CD28 beads in unstimulated and stimulated conditions, in the presence or absence of 5-AZA-CdR over a span of 12 days. n=8 per group, mean ± SEM. **B)** Cell cycle progression after 5 days of expansion. Each quadrant is one individual donor (n=4). **C)** Proportion of live versus dead cells after stimulation in the presence or absence of 5-AZA-CdR after 5 days of expansion. Each dot is one donor. ^*^p<0.05, paired t-test, two-sided, mean ± SD. **D, F, H**) Flow cytometry analyses of PD1 (**D**), LAG3 (**F**), and TIM3 (**H**) of one representative donor after 5 days of expansion. **E, G, I)** Flow cytometry analyses of PD1 (**E**), LAG3 (**G**), and TIM3 (**I**) of multiple donors. Each dot is one donor. ^*^p<0.05, ^**^p<0.01, ^***^p<0.001, n.s = non-significant, paired t-test, two-sided, mean ± SEM.

### DNMTi treatment during CD8+ T cell activation *ex vivo* modulates the expression of immune checkpoint and activation markers

Simultaneously, we examined expression of immune checkpoints receptors (PD1, LAG3 and TIM3) by flow cytometry. We observed increased expression of PD1 and LAG3, but reduced expression of TIM3 (Figures 4D-I), suggesting increased CD8+ T cell activation upon DNMTi treatment. These findings of increased activation were also reflected in significantly greater proportions of CD3^+^CD69^+^ cells (Figures S4B-C) and increased expression of the activation markers HLA-DR and CD25 (Figures S4D-E). However, while PD-1 and LAG3 are markers of T cell activation, they also have immune inhibitory functions and could suggest an exhaustion-like phenotype. In order to investigate whether the effector functions of CD8+ T cells were preserved or enhanced in the presence of DNMTi, we performed flow cytometric quantification for IFN-ɣ, TNF--α, and Granzyme B. As expected from *in silico* analysis and from the functional killing assays, we again found evidence for significant up-regulation in DNMTi treated cells, with elevated percentages of IFN-ɣ+ and TNF-α+ cells and higher Granzyme B MFI values (Figures 5A-F).

**Figure 5.**
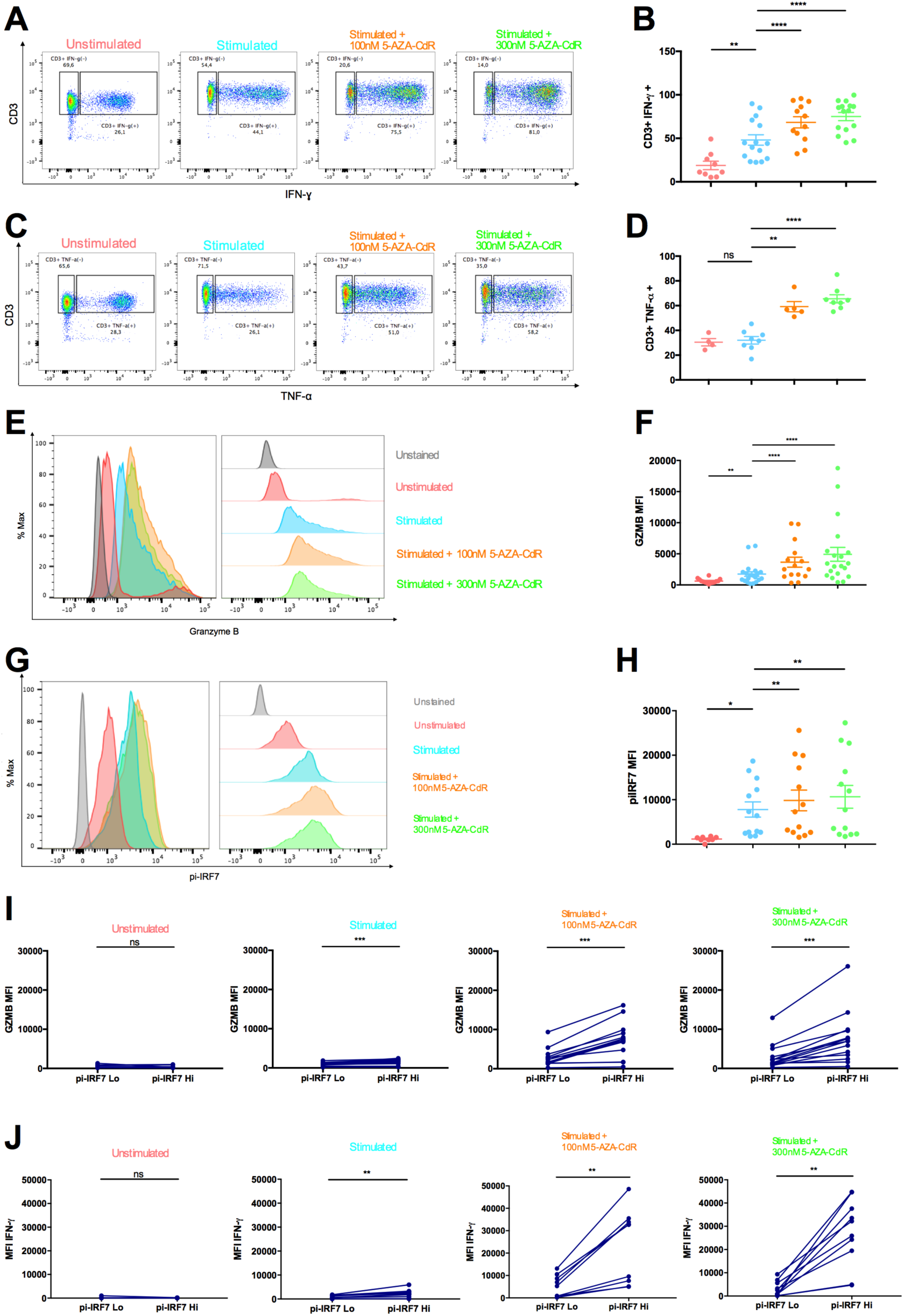
DNMTi treatment increases effector cytokines and proteolytic effector molecules through an IRF-7 dependent mechanism in activated human CD8+ T cells. **A, C)** Flow cytometry gating strategy to measure percent of CD3^+^IFN-ɣ^+^ T cells (**A**) and CD3^+^TNF-α^+^ (**C**) after *ex vivo* stimulation (Day 5) of CD8+ T cells in the presence or absence of 5-AZA-CdR in one representative donor. **B, D**) Combined percentages of CD3^+^IFN-ɣ^+^ T cells (**B**) and CD3^+^TNF-α^+^ (**D**) after *ex vivo* stimulation (Day 5) of CD8+ T cells in the presence or absence of 5-AZA-CdR in multiple donors. Each dot is one donor. ^*^p<0.05, ^**^p<0.01, paired t-test, two sided, mean ± SEM. **E-G**) Mean Fluorescence Intensity (MFI) of Granzyme B (**E**) and phospho-IRF7 (**G**) in one representative donor. **F, H)** Granzyme B (**F**) and phospho-IRF7 (**H**) MFI in multiple donors. Each dot is one donor. ^*^p<0.05, ^**^p<0.01, paired t-test, two sided, mean ± SD. **I-J)** Granzyme B MFI (**I**) and IFN-ɣMFI (**J**) in CD8+ T cell populations with phospho-IRF7 low versus phospho-IRF7 high in multiple donors. Each dot is one donor. ^**^p<0.01, ^***^p<0.001, n.s. = non-significant, paired t-test, two sided, mean ± SEM.

Additionally, we measured effector cytokine secretion by CD8+ T cells upon 5-AZA-CdR treatment through ELISA (Figures S5A-B), and measured mRNA gene expression of IFN-ɣ, TNF-α, and Granzyme B (Figures S5C-E). In agreement with our flow cytometric analyses, DNMTi treated CD8+ T cells expressed and secreted higher levels of these effector molecules.

### UpFregulation of effector markers is directly associated with IRF7 activation

We then evaluated whether the CD8+ T cells activation phenotypes upon DNMTi treatment were directly associated with IRF7 activation. Through flow cytometry, we measured levels of phosphorylated-IRF7. Indeed, we demonstrate that DNMTi treatment increased the levels of phosphorylated-IRF7 in stimulated CD8+ T cells (Figure 5G-H), agreeing with our IRF7 western blot analysis (Figure 3F).

Remarkably, when we sorted lower-quartile versus upper-quartile percentages of phospho-IRF7 CD8+ T cells (Figure S5F), we observed that Granzyme B and IFN-ɣ levels were significantly higher in the upper-quartile phospho-IRF7 cells, especially in DNMTi-treated cells (Figure 5I-J). Finally, we observed that increased Granzyme B levels in CD8+ T cells after DNMTi treatment was likely a consequence of IRF7 activation, as treatment with Type I Interferon alone can mimic the effects of 5-AZA-CdR treatment (Figure S5G-H). This suggests that the increase in these effector molecules after DNMTi treatment was likely a consequence of IRF7 activation, further indicating that the establishment of a viral mimicry state was responsible for these altered effector cytokine profiles.

## Discussion

Here, by leveraging a very large DNA methylation and gene expression dataset from TCGA across multiple tumor types, we show that IRF7 activation defines tumors marked by high levels of effector CD8+ T lymphocyte infiltration and elevated cytolytic activity. Notably, IRF7 activation itself is an independent predictor of cytolytic activity, and does not appear to be directly associated with neoantigen burden. Altogether, our data highlights that IRF7-a high tumors have an immune-reactive tumor microenvironment that is consistent with increased sensitivity to immune checkpoint inhibitors.

Furthermore, we identified that the increased activation and cytolytic activity in IRF7-high tumors is likely mediated by IRF7 activation directly in the immune cells. Building upon previous discoveries that DNA demethylating agents, at clinically relevant low doses, can activate IRF7 in tumor cells by inducing viral mimicry^1^, we report that DNMTi can also activate IRF7 and its downstream genes in CD8+ effector T cells and this can enhance T cell infiltration, activation and cytolytic activity. Mechanistically, this process is marked by DNA demethylation and up-regulation of repetitive elements, chiefly the LTR12 family of HERV9, activation of the dsRNA sensing pathway, leading to MAVS aggregation, IRF7 activation and up-regulation of the IRF7 associated transcriptional program. In total, our data indicate that DNMTi can exert significant anti-tumor activity by modulating CD8+ infiltration and effector function of individual CD8+ T cells, highlighting a pharmacomodulatory strategy to convert poorly infiltrated, IRF7-low “cold” tumors to infiltrated, IRF7-activated “hot” tumors (Figure 6). Moreover, our results agree with previously published data showing that the IRF7/IFNβ pathway is required for optimal anti-tumor activity of engineered CAR-T cells^28^.

**Figure 6.**
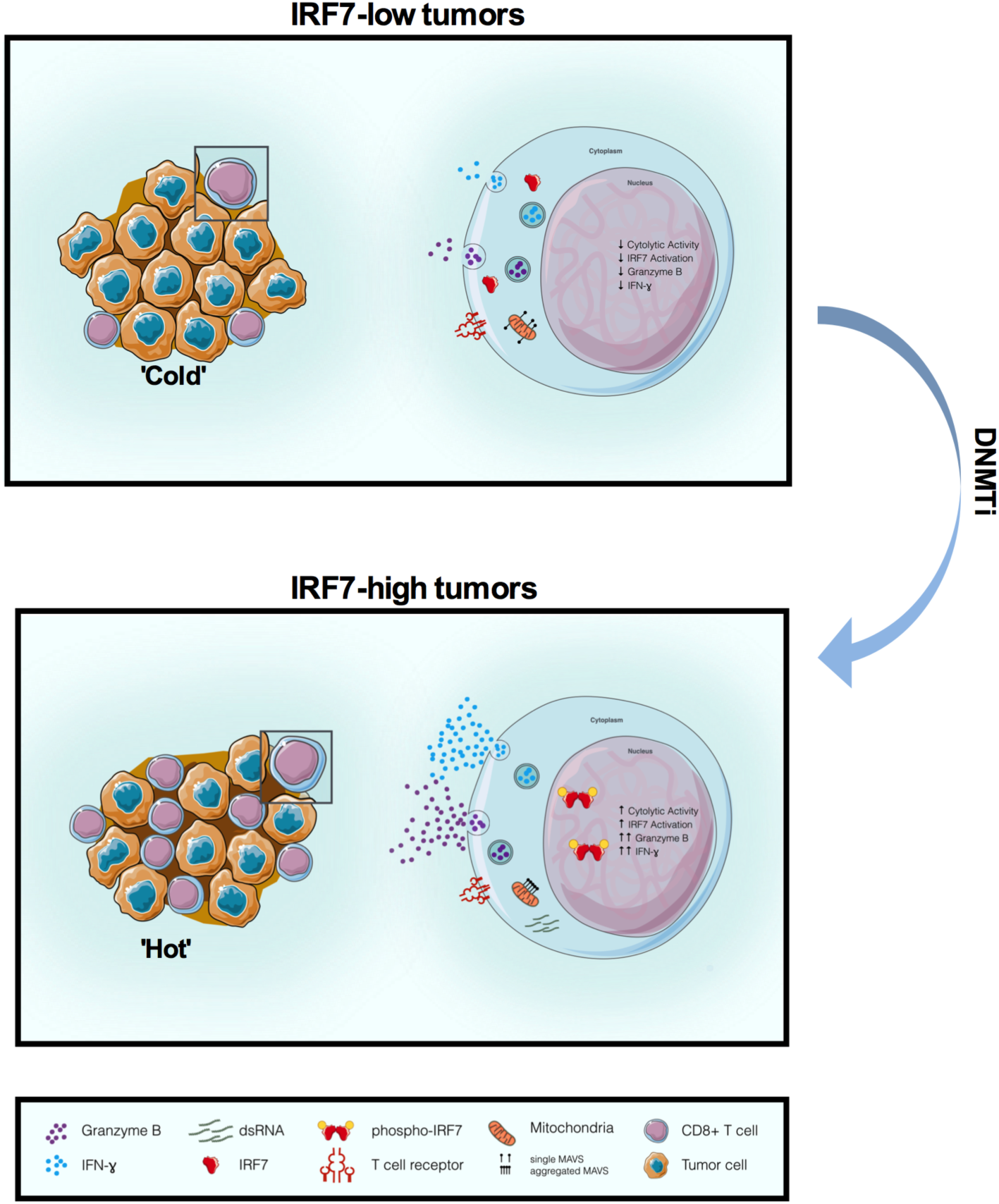
Schematic representation of how DNA demethylation drugs enhance CD8+ T cell activation.

Altogether, we have uncovered another layer of immune modulation by DNA demethylating drugs by directly increasing cytolytic activity of CD8+ T cells. Together with previous observations that global DNA demethylation induces viral mimicry in cancer cells and consequently increases tumor immunogenicity^1-3^, and recent data that DNA demethylating drugs can revert exhaustion-associated de novo methylation programs^29^, our data on CD8+ T cells activation suggests that DNA demethylating drugs can be potent immune-modulatory agents and sets the stage for clinical investigations combining DNMT inhibitors with immunotherapy strategies such as immune checkpoint blockade, bi-specific antibodies, adoptive cell therapy, or CAR-T cells.

## Methods

### Estimation of infiltrate composition from 450k methylation data

Illumina 450k array data were derived from SAGE synapse for multiple tumor types (Project ID: syn2812961) and processed using the MethylCIBERSORT suite of R functions^6^ to yield feature matrices for deconvolution. Missing probes were removed for eight cell types, and cancer-type specific cell line methylome profiles were compiled together during the development of MethylCIBERSORT, using the parameters AllGap = 0.3, FDR = 0.01 and nnGap = 0.1, and deconvolution was performed using the CIBERSORT server at cibersort.stanford.edu. For quality control, we imposed a correlation cutoff of at least 0.85 between imputed and original beta-values. This produced a set of 6913 tumors with estimates of infiltrate composition.

### Definition of clusters based on IRF7 genes

RNA-seq data were obtained from SAGE Synapse (Project ID: syn4301332) for 7769 tumors in the form of aggregated level 3 RSEM expression estimates, which were transformed to log2 counts per million and then quantile normalized. An integrated dataset of 5,996 tumors, across 19 tumor types, was then generated by reducing samples to those that had both matched RNA-seq data and infiltrating cell type estimates. The *pamk* function from the *fpc* package was used to partition the samples in the integrated dataset into distinct clusters, with the optimal number of clusters picked from a range of 2-10 using average silhouette width as the selection criterion. Associations between IRF7 signature clusters and estimated abundances of individual infiltrating cell types were tested using Wilcoxon’s Rank Sum Test with Benjamini-Hochberg correction for multiple testing.

### Modelling,cytolytic activity and link with IRF7 activation profile

Cytolytic activity (CYT) was defined as the geometric mean of PRF1 and GZMA expression as previously described^11,30^. To measure the influence of IRF7 activation on the activity of the infiltrate, a linear model was fit regressing log2(CYT) with different immune cell populations and IRF7 cluster as explanatory variables. Univariate associations between IRF7 cluster and cytolytic activity were tested using Wilcoxon’s Rank Sum Test with Benjamini-Hochberg correction for multiple testing. We also defined an IRF7 activation score using ssGSEA^14^, performed using the GSVA Bioconductor package^31^, for further regression modelling.

### Modelling correlates of IRF7 activation clusters

Neoantigen estimates were obtained for samples from The Cancer Immunome Atlas (https://tcia.at) and Wilcoxon’s Rank Sum Tests with Benjamini-Hochberg FDR correction for multiple testing were used to determine association with IRF7 cluster. The transcriptional correlates of IRF7 cluster membership were identified by fitting a *limma*-trend^32^ model with correction for mean-variance relationships during empirical Bayes estimation. DEGs were defined at 2 fold-change, BHFDR < 0.001.

### Modelling the origins of the IRF7 activation signal

We used ssGSEA scores for the IRF7 signature to test for correlations with estimated abundances of different cell types and purity per cancer type using Spearman’s Rank Correlations with Benjamini-Hochberg correction for multiple testing. ssGSEA scores for the IRF7 signature were then regressed against estimates of infiltrating cell types to evaluate the contribution of different factors to the overall ssGSEA enrichment signal. For validation, flow-sorted populations from colorectal cancers were obtained from GSE39397. Affymetrix data were rma-normalized and then collapsed to genewise-values using mean-summarization. ssGSEA scores were then computed using the GSVA R package. Paired Wilcoxon’s Tests were used to compare enrichment scores in leukocyte fractions with their matched tumor fractions.

### Analysis of single cell data

We obtained SMART-Seq2 counts for dissociated single cells from melanomas from GSE72056^15^. The data were normalized using the scran package^33^ and for analysis of TILs, we selected all cells defined as T cells in the original paper. Consensus PAM clustering was then used to identify subpopulations defined by CD4/CD8+A/CD8+B expression. ssGSEA scores and CYT were computed as described and linear regression, Spearman’s correlations or Wilcoxon’s Rank Sum tests were used for hypothesis testing.

### Analysis of transcriptional influences of 5-AZA-CdR treatment on CD8+ T cells

mRNA for RNA-seq was extracted using the RNeasy Plus Mini Kit (Qiagen, cat # 74104). mRNA was isolated from cells harvested from unstimulated T cells collected at Day 0, and T cells stimulated in the presence or absence of 300nM 5-AZA-CdR at Day 5 post-stimulation. RNA sample quality was measured by Qubit (Life Technologies) for concentration and by Agilent Bioananlyzer for RNA integrity. All samples had RIN above 9. Libraries were prepared using the TruSeq Stranded mRNA kit (Illumina). Final cDNA libraries were verified by the Agilent Bioanalyzer for size and concentration quantified by qPCR. The libraries were sequenced by the Princess Margaret Genomics Centre on an Illumina HiSeq 2000 as a pair-end 100 cycle sequencing run using v3 reagents to achieve a minimum of ∼30 million reads per sample.

Fastq files were pseudoaligned to GENCODE hg19 lncRNAs and protein-coding genes and quantified using *Kallisto*. The reads were parsed into R with a custom function. Low expressed reads (< 0.5 counts per million on average) were removed before voom precision weight computation on quantile-normalized data. Linear models were fit using limma with treatment status and donors as a blocking factor and differentially expressed genes were defined between 5-AZA-CdR-treated and untreated stimulated CD8+ T cells at Day 5 at FDR < 0.05, 2 fold-change. Significance of enrichment for IRF7 cluster-associated DEGs within Aza-associated DEGs in the model system was computed by sampling 10,000 random sets of 1043 genes (the size of the 5-AZA-CdR signature) and computing the frequency of an observed overlap at least as great as observed with IRF7 cluster-associated DEGs.

### Analysis of methylome changes in 5-AZA-CdR treated CD8+ T cells

RRBS was carried out following the protocol indicated previously^34^, with some minor modifications. In brief, 25 ng of DNA was library prepared (end-repaired, A-tailed and adapter-ligated with Illumina methylated barcodes) prior to bisulfite conversion using Zymo EZ DNA Methylation Kit (cat# D5001), following the Illumina protocol in the kit. The bisulfite converted libraries were then gel size selected for fragments between 160bp to 400bp in size, prior to amplification and sequencing. The samples were sequenced by the Princess Margaret Genomics Centre, using an Illumina HiSeq 2000, single-read, v4 chemistry, with a minimum of 40 million reads per sample.

Trim Galore was used to quality-trim (Phred score cutoff 20) and then adapter trim fastq files. Reads greater than 20bp in length were retained and processed with Bismark as RRBS data. A feature by sample matrix of Beta-values was defined for all CpGs covered by at least 10 reads in all samples.

MVPs were computed using a custom wrapper for limma-based functions with correction for mean-variance trend and were defined at absolute Delta-Beta > 0.3, FDR < 0.01. Expression data were then integrated with DMR and MVP calls by filtering for concordant associations (enhancer, intergenic repeat sequence CpGs and promoter hypermethylation inversely associated with DEGs, body methylation directly associated with DEGs) for downstream annotation and analysis.

### CD8+ T cell isolation and culture

Human CD8+ T cells were isolated using a negative selection kit (Miltenyi Biotec USA, Cat# 130-096-495) from peripheral blood mononuclear cells (PMBC) collected from healthy blood donors. Blood from healthy donors was collected in the clinic after obtaining appropriate informed consent under an UHN-REB (#11-0343-CE) approved protocol. At least 92% purity was obtained using this kit.

Purified CD8+ T cells were seeded in specialized non-adherent round-bottom 96-well plates at a concentration of 10,000 cells per well, in a final media volume of 200uL. The media used for CD8+ T cell cultures was RPMI 1640 conditioned with 2mM HEPES, 0.1% Pen Strep 0.1% L-glutamine, 10% Human Serum, and 10IU/mL human recombinant IL-2.

Treatments included: unstimulated (no beads added); stimulated; stimulated plus 100nM 5-AZA-CdR; and stimulated plus 300nM 5-AZA-CdR. In order to stimulate CD8+ T cells, anti-CD3/CD28 beads (ThermoFisher Scientific, Cat#11131D) were added to the wells at a 1 bead: 2 cells ratio and left in the wells until the last time point of data collection (Day 12). For the stimulation of CD8+ T cells with 5-AZA-CdR (Sigma-Aldrich, Cat#A3656), two concentrations were used during the CD8+ T cell expansion protocol: 100nM and 300nM. Due to the half-life of 5-AZA-CdR, fresh media was replaced every 48 hours (from Day 0 onwards) to maintain respective concentrations of 5-AZA-CdR.

In the conditions where Type I IFNs was added during stimulation, 10ng/mL of IFN-α (cytokine) was used. Fresh media was replaced every 48 hours (from Day 0 onwards) to maintain concentrations of IFN-α.

### Cell count and cell cycle assay analysis

CD8+ T cell numbers from all conditions were counted using the automated cell-counting machine Vi-CELL XR (Beckman Coulter). In brief, cell count was performed at Days 2, 4, 6, 8, 10 and 12 post-stimulation. Prior to counting, anti-CD3/CD28 beads were removed using a DynaMag-2 Magnet (ThermoFisher Scientific, Cat#12321D).

Prior to CD8+ T cell stimulation with anti-CD3/CD28 beads and 5-AZA-CdR treatments, a portion of CD8+ T cells were stained with CellTrace™ Violet Cell (CTV) Proliferation dye (ThermoFisher Scientific, Cat#C34557) to measure cell cycle progression at Day 5 post-stimulation. 5-AZA-CdR is only incorporated during DNA synthesis during cell division; hence, by Day 5, three doses of 5-AZA-CdR had been administered *in vitro* to ensure thorough demethylation. At Day 5 post-stimulation, cells were harvested, washed with FACS washing buffer (PBS + 0.5% BSA). Cell cycle progression was measured through flow cytometry using the BD FACS Canto™ II system.

### Semidenaturing Detergent Agarose Gel Electrophoresis

MAVS aggregation assay was performed following a previously published semidenaturing detergent agarose gel electrophoresis (SDD-AGE) (Alberti et al., 2009). Mitochondrial proteins were purified using the Qproteome Mitochondria Isolation Kit (Qiagen, cat#37612). Equal amounts of mitochondrial extract were run in a vertical 1% agarose gel containing 0.01% SDS at constant 50V for 90 minutes. Following that, proteins were immunoblotted to a PVDF membrane using the Trans-Blot Turbo Transfer System (Biorad). Lastly, the membrane was blotted with the anti-MAVS antibody (Abcam, ab31334) and developed accordingly.

### Flow cytometry analysis

Fluorescence readings were performed using the BD FACS Canto™ II system. For flow cytometric analyses, CD8+ T cells were stained with a cell viability dye, LIVE/DEAD^®^ Fixable Violet Dead Cell Dye (ThermoFisher Scientific, Cat#L-34963). Next, extracellular surface markers were stained at 4°C for 30 minutes, and cells were fixed with Fixation Buffer 4°C for 30 minutes (eBioscience, Cat#00-5523-00). Cells were washed 2X using FACS washing buffer prior to analysis using the BD FACS Canto™ II system. Results and histograms were generated using FlowJo software.

For intracellular marker staining of functional cytokines, CD8+ T cells were further stimulated with Cell Stimulation plus Protein secretion inhibitor Cocktail (Affymetrix, Cat#00-4975-03) for 5 hours. Afterwards, cells were stained with cell viability dye and cell surface markers as described above, and CD8+ T cells were washed twice with Permeabilizing Buffer (eBioscience, Cat#00-5523-00). Intracellular antibody staining was performed in Permeabilizing Buffer at 4°C for 30 minutes. Cells were washed twice prior to analysis with BD FACS Canto™ II system. For the staining of phospho-IRF7, Fix Buffer I (BD Bioscience, Cat#557870) and Perm Buffer III (BD Bioscience, Cat#558050) were used for fixation and permeabilization of cells, respectively. For the staining of Annexin V and Propidium Iodide (PI), Annexin V Binding Buffer was used (ThermoFisher Scientific, Cat#V13246). Results and histograms were generated using FlowJo software.

Fluorescence minus one (FMO) was used as the unstained control for measuring extracellular markers; isotype control was used as the unstained control for measuring intracellular markers. Antibodies used: anti-human CD69 (Biolegend), anti-human CD3 (Biolegend, BD Pharmingen), anti-human/mouse Granzyme B (Biolegend), anti-human TNF-α (Biolegend), anti-human IFN-ɣ (Biolegend), anti-human CD366 (TIM-3) (Biolegend), anti-human CD279 (PD-1) (Biolegend), anti-human CD223 (LAG3) (eBioscience), anti-human phospho-IRF7 (BD Bioscience), Annexin V (Biolegend), Propidium Iodide Staining Solution (BD Pharmingen). The antibodies’ catalog numbers are available in Supplementary Table S11.

### RT-qPCR

To measure mRNA levels using reverse transcription polymerase qualitative chain reaction (RT-qPCR), mRNA was isolated from cells harvested from unstimulated T cells collected at Day 0, and T cells stimulated in the presence or absence of 300nM 5-AZA-CdR at Days 1, 2, 3, 5, and 7 post-stimulation. mRNA was extracted using RNeasy Plus Mini Kit (Qiagen, Cat#74104).

To perform the reverse transcription, 200 ng of mRNA were used for conversion to cDNA. The recommended protocol from the SuperScript III First-Strand Synthesis System (ThermoFisher Scientific, Cat#18080051) was followed. Converted cDNA was diluted 1:50 prior to qPCR thermal cycling to amplify gene of interest based of the primer used. Next, the SYBR Select Master Mix (ThermoFisher Scientific, Cat#4472903) protocol was performed as recommended. The thermal cycling for gene amplification was as follows: 95°C for 10 minutes, 40 cycles of 95°C for 15 seconds, 60°C for 1 minute. A melting curve analysis was performed to confirm no genomic contamination or primer dimer products. Each transcript level was normalized by the acidic ribosomal phosphoprotein P0 (RPLP0) housekeeping gene. To analyze gene amplification, a fold-change calculation was performed based on Cycle Threshold (CT) value normalized against RPLP0 levels, followed by normalization of the conditions stimulated in the presence or absence of 300nM 5-AZA-CdR against the unstimulated condition.

Primers: Granzyme B (AGATGCAACCAATCCTGCTT; CATGTCCCCCGATGATCT), IFN-ɣ (GAGTGTGGAGACCATCAAGGA; CTGTTTTAGCTGCTGGCGAC), TNF-α (GACAAGCCTGTAGCCCATGT; TCTCAGCTCCACGCCATT), RPLP0 (CAGACAGACACTGGCAACA; ACATCTCCCCCTTCTCCTT).

### ELISA

To measure the cytokine levels found in the supernatant, an enzyme-linked immunosorbent assay (ELISA) was performed. Supernatant from cell cultures were collected at Days 2, 4, and 6 post-stimulation to measure TNF-α and IFN-ɣ cytokine production. ELISA kits were purchased from eBioscience (Human IFN-ɣ ELISA Ready-Set-Go, Cat#88-7316; Human TNF-α ELISA Ready-Set-Go, Cat#88-7346), and the manufacturer’s protocol was followed.

### *in vitro* killing assay using T cell bi-specific (TCB) antibodies

Generation and maintenance of MCF10A cells stably expressing HER2 and p95HER2 and PDX118, a HER2-positive primary breast cancer cell line derived from a patient-derived xenograft, have been previously described (Parra-Palau JL et al, J Natl Cancer Inst. 2014 Sep 24;106(11). pii: dju291. doi: 10.1093/jnci/dju291). Cells were labeled with CellTrace CFSE according to manufacturer’s protocol (ThermoFisher).

Peripheral blood mononuclear cells (PBMCs) were obtained by Ficoll-Paque Plus density gradient centrifugation of buffy coats. Subsequently, CD8+ T cells were isolated using the human CD8+ T Cell Isolation Kit (Miltenyi Biotec) following manufacturer’s instructions. CD8+ cells were stimulated with Dynabeads Human T-Activator CD3/CD28 in presence of 10 IU/mL of IL-2 for 5 days and, when indicated, 100nM Decitabine. Medium was replenished every 2 days. After 5 days, cells were harvested and beads removed before cell counting.

CFSE-labeled MCF10A and PDX118 were co-cultured at 1:10 ratio for 16 hours and 1:5 ratio for 24 hours, respectively, with CD8+ T cells. HER2-TCB was added at 1nM or 0.5nM for MCF10A or PDX118 co-cultures, respectively. CFSE-positive cell number was evaluated by flow cytometry using BD LSRFortessa cell analyzer (BD Bioscience).

### OT-1 TCR-specific model

OT-1 mice expressing the OVA_257–264_/K^b^-specific TCR were bred in-house. All work with mice was in compliance with the guidelines of the Institutional Animal Use and Care Committee.

### *In vitro* cytotoxic killing assay using OT-1 specific T cells

Activated OT-1 T cells were generated *in vitro* in the presence of Concanavalin A and IL-2 to be used as effector cells in the *in vitro* cytotoxic assay. Briefly, OT-1 mouse spleen was homogenized and red blood cells were lysed for 2 min with 2 mL of ACK buffer buffer (0.15 M NH_4_Cl, 1 mM KHCO_3_, and 0.1 mM EDTA). Single cell suspension of OT1 splenocytes were cultured in complete media (RPMI 1640 containing 10% heat-inactivated FBS, 100μM 2-mercaptoethanol, 1% L-glutamine, 1 mM sodium pyruvate, 1% penicillin/streptomycin, 100μM MEM NEAAS, 50ng/mL Gentamicin) with 5 μg/mL of Concanavalin A in the presence or not of 0.3μM of 5-AZA-CdR and incubated at 37 °C and 5% CO_2_ for 48 hours. Then, cells were harvested, washed and cultured in complete media with 10 U/mL of IL-2 and in the presence or not of 0.3μM of 5-AZA-CdR. After 48 hours, the same concentration of IL-2 was added to each well following an expansion at Day 6. On the next day, viable lymphocytes were recovered by centrifugation on Ficoll gradient at 500 *g* for 30 min and used as effector cells. For target cells, RMA cells were labeled with carboxyfluorescein diacetate succinimidyl diester (CFSE; Molecular probes, Eugene, Oregon, USA) at two concentrations: 10μM (CFSE_high_) and 1μM (CFSE_low_). Fluorescence-labeled cells were either pulsed or not with 10 μM of OVA peptide SIINFEKL H-2K_b_ (257-264) (InvivoGen, San Diego, USA) making up two populations that were mixed together in equal proportion (1:1). Then, 2×10^4^ of target cells were plated in a 96 round-bottom well plate together with effector cells in a 1:1 proportion (target cells/effector cells). After 6 hours, cells were harvested, washed and fixed in 1% paraformaldehyde following analysis in flow cytometry (BD FACSCanto II; BD Biosciences, San Jose, CA). A deviation from the 1:1 ratio of control and target cell populations indicated antigen-specific lysis of target cells.

### *in vivo* CT26 tumor model

8-weeks-old females BALB/c mice were obtained from Charles River Laboratories and housed at the CRCHUM animal facility. Animals (10 per group) were injected subcutaneously with 500,000 CT26 cells in 100μ L PBS. Mice were weighed on Day 4 and treated intraperitoneally with 0.5 mg/kg of Decitabine (Sigma; A3656) for 5 consecutive days starting on Day 5 after tumor inoculation. CD8+ T cells depletion was performed on Day 4, 5 and 11 with anti-CD8+ clone 53-5.8 antibody (BioXCell; 100 μg/host) respectively. CD8+ T cell depletion was confirmed by flow cytometry. Tumor size (mm2) was monitored every 2-3 days. Mice were sacrificed on Day 19 and tumors were harvested in OCT media and stored at -80°C.

## Statistical Analysis

Flow cytometry, ELISA, RT-PCR gene expression assays were assessed using the software GraphPad Prism 7. Specific test used are described in figure legend.

## Supplementary Figures

**Figure S1.**
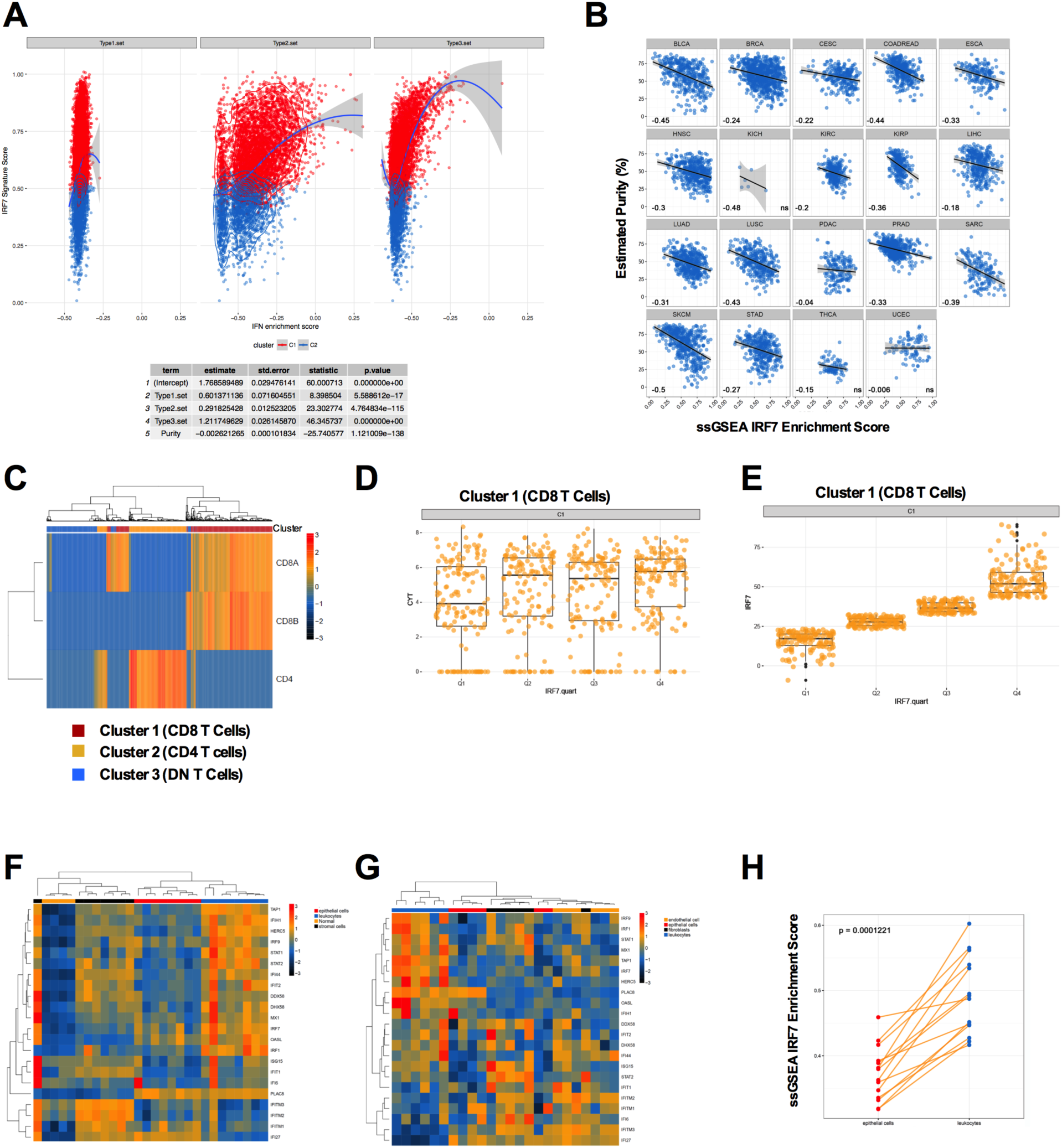
A) Comparison of IRF7 ssGSEA enrichment score (Y-axis) versus interferon gene enrichment score (X-axis), with the table denoting effect sizes and significance statistics from multivariate linear modelling of IRF7 score as a function of interferon gene set scores **B)** Correlation scatterplots of IRF7-signature ssGSEA scores (X-axis) versus estimated tumor purity (Y-axis) across multiple tumor types, values represent Spearman’s Rho, with ns = not significant at FDR > 0.05. **C**) Heatmap showing consensus PAM clustering of TIL data from Tirosh et al highlighting patterns of CD8+ subunit and CD4 expression **D)** Boxplots showing CYT by quartile of IRF7 enrichment score in CD8+ TILs from Tirosh et al. **E)** Boxplots showing distribution of IRF7 enrichment score by score quartile in CD8+ TILs from Tirosh et al **F-G)** Heatmaps of the expression of IRF7 signature genes in microarray analyses of flow sorted subpopulations of colorectal cancers from GSE39397, on two distinct microarray platforms. Rows represent expression Z-scores for each gene, columns represent samples. **H)** Sorted leukocyte fractions from GSE39397 show significantly higher ssGSEA enrichment scores for IRF7 activation relative to epithelial fractions from matched tumors in GSE39397. P-value from Paired Wilcoxon’s Rank Sum Test.

**Figure S2.**
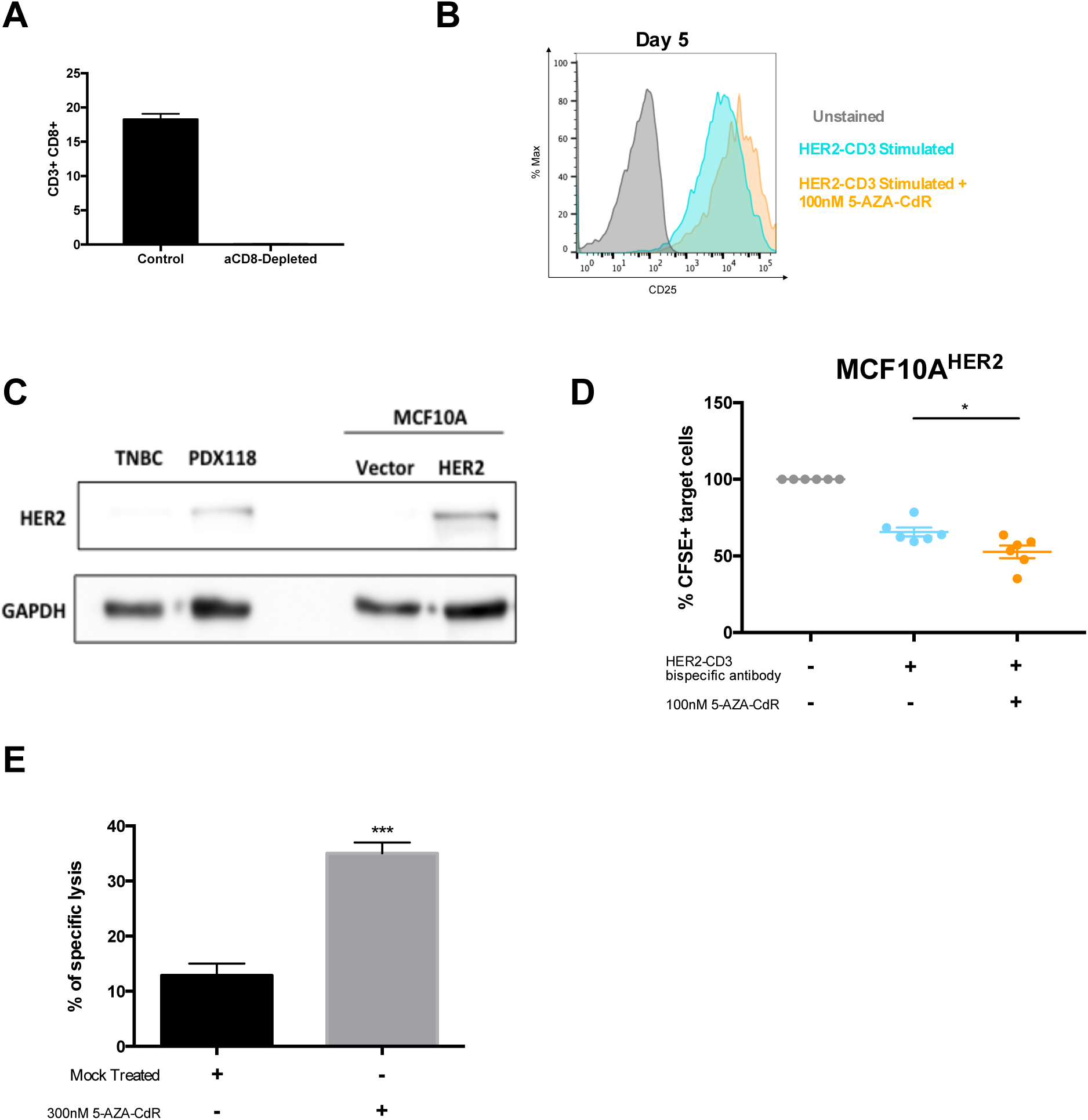
A) Percent of live CD3+CD8+ T cells before and after anti-CD8+ antibody depletion. **B)** Flow cytometry analyses of CD25 in human CD8+ T cells unstained (grey), stimulated with the HER2-CD3 bispecific antibody (Blue) and stimulated with the HER2-CD3 bispecific antibody plus 100uM 5-AZA-CdR treatment (orange). **C)** Left: Anti-HER2 western blots for the breast cancer patient-derived HER2 cells used in Figure 2F (PDX118) and a triple negative breast cancer (TNBC) sample used as a negative control. Right: Anti-HER2 western blots for the non-transformed MCF10A mammary epithelial cell transfected with an empty vector or a HER2 vector. **D)** Quantification of surviving target cells (MCF10A^HER2^) after the killing assay using human CD8+ T cells treated or not with DNMTi in the presence of anti-HER2-CD3 bispecific antibody. ^*^p<0.05 versus the untreated group. Paired t-test, mean ± SEM). **E)** Killing assay of OVA pulsed target cells by CD8+ T cells from OT-I mice *ex vivo* treated with DNMTi.

**Figure S3.**
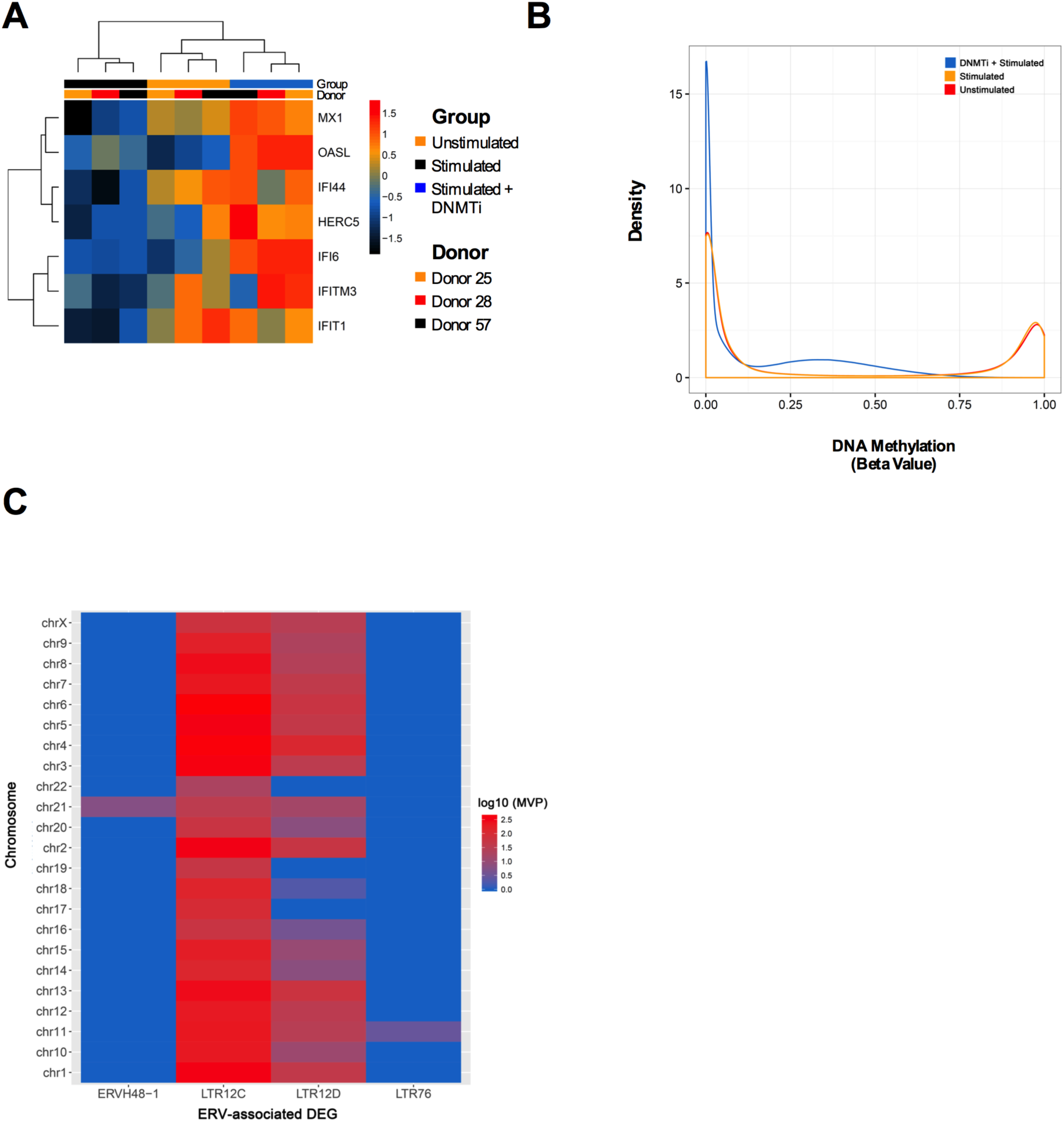
A) Heatmap showing the IRF7 signature genes that are differentially expressed (FC > 2, FDR < 0.05) in activated CD8+ T cells in the presence of 300nM 5-AZA-CdR versus activated CD8+ T cells without DNMTi treatment. Rows represent expression Z-scores, columns represent samples. **B)** Density plots of average beta-values by group. DNMTi induces global hypomethylation in stimulated CD8+ cells relative to unstimulated and mock-treated stimulated cells. **C)** Heatmap of DNA demethylated repeats associated with ERV-derived DEGs across chromosomes. Y-axis = chromosome, X-axis = ERV, colour = log10(number of MVPs on that chromosome associated with the repeat)

**Figure S4.**
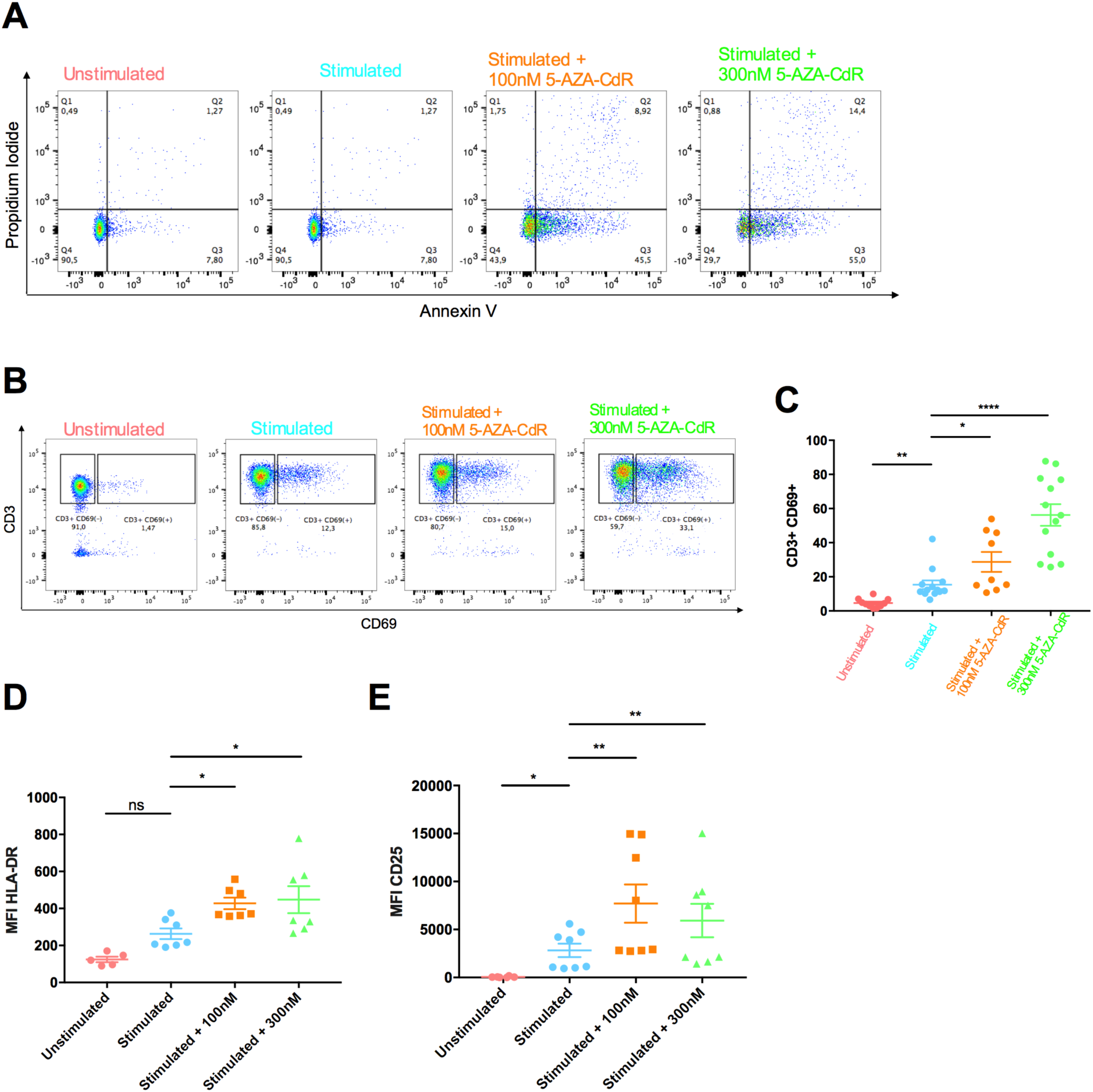
A) Measurements of apoptosis markers Annexin V and Propidium Iodide in CD8+ T cells after stimulation in the presence or absence 5-AZA-CdR in one representative donor. **B)** Flow cytometry gating strategy to measure percent of CD3^+^CD69^+^ T cells after stimulation in the presence or absence 5-AZA-CdR in one representative donor. **C)** Combined percentages of of CD3^+^CD69^+^ T cells in multiple donors. Each dot is one donor. ^*^p<0.05, ^**^p<0.01,^****^p<0.0001, paired t-test, two sided, mean ± SEM. **D-E)** Mean Fluorescence Intensity (MFI) measurement of HLA-DR (**D**) and CD25 (**E**) expression by CD8+ T cells after 24 hours post anti-CD3 and anti-CD28 stimulation. Each dot is one donor. ^*^p<0.05, ^**^p<0.01, n.s. non-significant, paired t-test, two sided, mean ± SEM.

**Figure S5.**
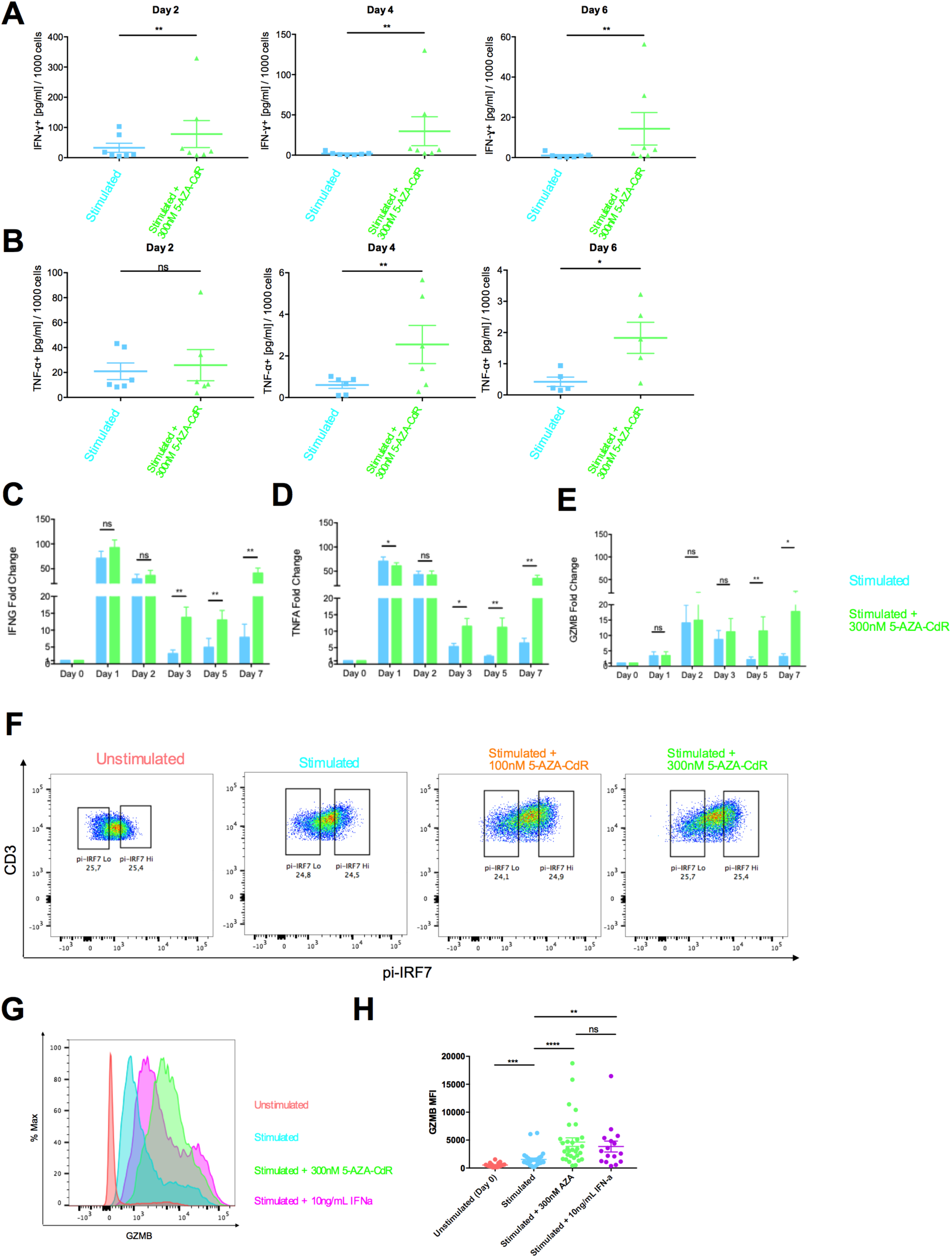
**A-B**)IFN-ɣ (**A**) and TNF-α (**B**) cytokine secretion by CD8+ T cells after stimulation in the presence or absence 5-AZA-CdR quantified by ELISA. Each dot is one donor. ^*^p<0.05, ^**^p<0.01, n.s. non-significant, ratio paired t-test, two sided, mean ± SEM. **C-E**) Gene expression profile of IFNG (**C**), TNFA (**D**), and GZMB (**E**) genes quantified by RT-qPCR. n=5. ^*^p<0.05, ^**^p<0.01, n.s. non-significant, ratio paired t-test, two sided, mean ± SEM. **F)** Flow cytometry gating strategy to sort out top quartile (phospho-IRF7^High^) versus bottom quartile (phospho-IRF7^low^) of activated-IRF7 CD8+ T cell populations after stimulation in the presence or absence 5-AZA-CdR in one representative donor. **G-H)** Granzyme B MFI in CD8+ T cells stimulated in the presence of DNMTi (light green), 10ng/mL Interferon-α (pink) or mock (light blue) in one representative donor (**G**) and in multiple donors (**H**). Each dot is one donor. ^**^p<0.01, ^****^p<0.0001, n.s. non-significant, paired t-test, two sided, mean ± SEM.

